# Genome-wide analyses of chromatin interactions after the loss of Pol I, Pol II and Pol III

**DOI:** 10.1101/2020.01.28.923995

**Authors:** Yongpeng Jiang, Jie Huang, Kehuan Lun, Boyuan Li, Haonan Zheng, Yuanjun Li, Rong Zhou, Wenjia Duan, Yuanqing Feng, Hong Yao, Cheng Li, Xiong Ji

**Author notes:** These authors contributed equally to this work (Yongpeng Jiang, Jie Huang, Kehuan Lun, Boyuan Li).

## Abstract

**Background:** The relationship between transcription and the 3D genome organization is one of the most important questions in molecular biology, but the roles of transcription in 3D chromatin remain controversial. Multiple groups showed that transcription affects global Cohesin binding and genome 3D structures. At the same time, several studies have indicated that transcription inhibition does not affect global chromatin interactions.

**Results:** Here, we provide the most comprehensive evidence to date to demonstrate that transcription plays a marginal role in organizing the 3D genome in mammalian cells: 1) degraded Pol I, Pol II and Pol III proteins in mESCs, and showed their loss results in little or no changes of global 3D chromatin structures for the first time; 2) selected RNA polymerases high abundance binding sites-associated interactions and found they still persist after the degradation; 3) generated higher resolution chromatin interaction maps and revealed that transcription inhibition mildly alters small loop domains; 4) identified Pol II bound but CTCF and Cohesin unbound loops and disclosed that they are largely resistant to transcription inhibition. Interestingly, Pol II depletion for a longer time significantly affects the chromatin accessibility and Cohesin occupancy, suggesting RNA polymerases are capable of affecting the 3D genome indirectly. So, the direct and indirect effects of transcription inhibition explain the previous confusing effects on the 3D genome.

**Conclusions:** We conclude that Pol I, Pol II, and Pol III loss only mildly alter chromatin interactions in mammalian cells, suggesting the 3D chromatin structures are pre-established and relatively stable.

## Background

The relationship between transcription and the 3D chromatin structures is one of the most fundamental questions in the post-genomic era [1–4]. Mounting evidence has shown that transcription activities correlate with more DNA interactions in development and diseases [5–8]. Topologically associating domains (TADs) boundaries or insulated neighborhoods or CTCF loop domains usually enrich active transcription with both protein machinery and noncoding RNAs[9–11]. In *D. melanogaster*, transcription predicts chromatin organization [12–14], suggesting a potential causal relationship between transcription and 3D chromatin landscape.

Recent studies combining knock out with inhibition showed that transcription could relocate Cohesin over mammalian chromatin [15], implicating that Pol II may regulate the 3D genome via its impact on Cohesin. Blocking of transcription elongation in mammalian cells increases Cohesin binding and loop formation at CTCF binding sites within the gene bodies [16]. Inhibit *D. melanogaster* transcription alters chromatin interactions both within and between domains [12–14]. However, it is unclear whether Pol II regulates 3D chromatin landscapes via Cohesin directly.

Many early development-related studies have revealed insights for the transcription and 3D chromatin landscape. Pol II transcription inhibition during the early development of mouse embryo does not affect global TAD structures [17, 18], while it is difficult to interpret because of the relatively low sequencing depth of these experiments and developmental arrest after transcription inhibition. Also, the chromatin organizations of transcriptionally inactive mature oocytes and sperms are quite similar to the embryonic stem cells [17, 19–21], implicating that it might not be transcription per se, but transcription-associated proteins contribute to the 3D genome organization.

The unchanged phenotype after transcription inhibition in mammalian cells is usually based on aggregate analyses of all chromatin loops [17, 18, 21–23]. It is hard to evaluate the contribution of transcription, as CTCF and Cohesin play a predominant role in the 3D chromatin landscape, and they occupy most of the loops in mammalian cells [24–26]. It is immature to conclude that transcription has no impact on chromatin interactions in mammals because one of the critical evidence is still missing in the field, which is to precisely evaluate the roles of Pol II in the 3D genome in the absence of CTCF and Cohesin.

RNA polymerases synthesize RNA, forming DNA, RNA, and protein ternary complexes in the nucleus [27–29]. Pol I, Pol II, and Pol III are three distinct DNA-dependent RNA polymerases that function together with thousands of transcriptional and chromatin regulators to synthesize rRNA, mRNA, and tRNA, respectively [9, 30–36]. Although the structure and function of RNA polymerases have been studied for 50 years [37], in particular, the roles of Pol I and Pol III in 3D chromatin organizations are poorly investigated.

To explore the function of RNA polymerase proteins in 3D chromatin organization, we specifically degraded the largest subunits of Pol I, Pol II, and Pol III in murine embryonic stem cells. Global chromatin interactions and higher resolution interactions among regulatory elements remain unchanged after Pol II depletion. Chromatin organization reforms during mitotic exit in the absence of Pol II, supporting the pre-established model for the 3D genome. We then identified Pol II bound but Cohesin and CTCF unbound loops, and they are largely resistant to transcription inhibition both in our and public datasets from different labs. Additionally, acute depletion of Pol I and Pol III also does not result in global and local changes for the 3D genome. These results collectively demonstrate that RNA polymerases play a marginal role in organizing 3D chromatin landscapes in mammalian cells. Interestingly, long-time depletion of Pol II decreases chromatin accessibility and Cohesin occupancy over chromatin. We propose that immediate transcription inhibition does not affect 3D chromatin structures, but the indirect effects of transcription inhibition do.

## Results

### Acute degradation of Pol I, Pol II, and Pol III in mESCs with auxin-inducible degron technology

To investigate the roles of RNA polymerase proteins in 3D chromatin organization, we subjected mESCs to RNA polymerases degradation with degron technology. RNA polymerase-mediated transcription could be inhibited with different inhibitors, while they either lost the capacity to distinguish different RNA polymerases or had specificity for RNA polymerase (such as alpha-amanitin) but worked at high concentrations and take a long time [38, 39]. The degron system was applied to RNA polymerase subunits because it degrades protein rapidly and accurately (Fig. 1a) [40, 41]. The auxin-inducible degron system was set up in murine embryonic stem cells: exogenously expressed OsTIR1 was inserted into the Rosa26 locus under the Tet-on promoter, and then the protein of interest was tagged at its C terminal with mAID-GFP by CRISPR genome editing followed by single clone selection (Fig. S1a). The mAID tag could be recognized and degraded by OsTIR1 in the presence of auxin (Fig. 1a). The C-terminal domain of the largest subunit of RNA polymerase was tagged with mAID-GFP. This system allows us to specifically investigate endogenous Pol I (Rpa1), Pol II (Rpb1), and Pol III (Rpc1) with the GFP tag and the phenotype for loss of function with the mAID tag.

**Fig. 1.**
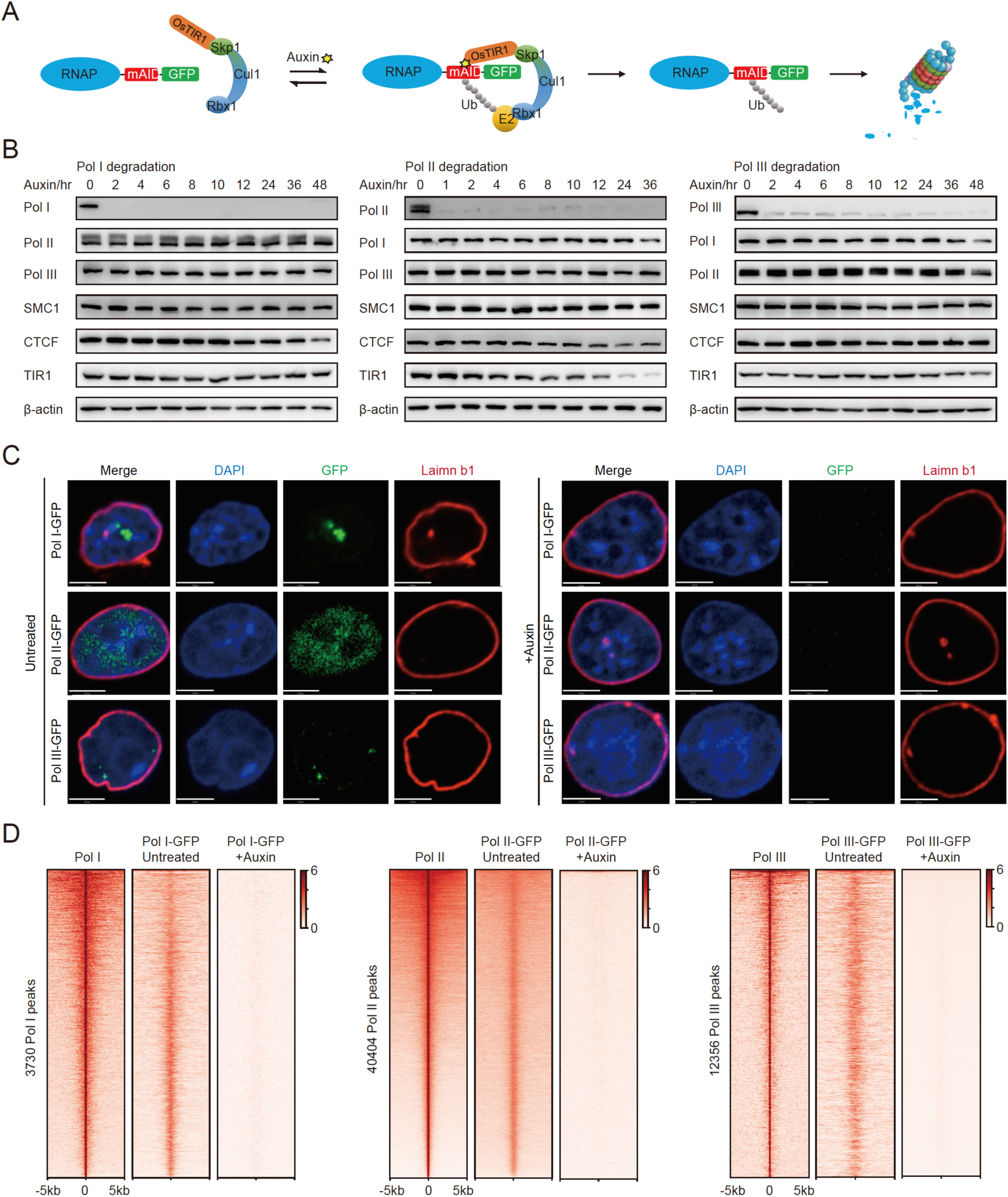
Rapid depletion of endogenous Pol I, Pol II, and Pol III proteins in mESCs. a) Schematic of auxin-inducible degron technology. Endogenous RNA polymerases were fused to the mAID-GFP tag in the C terminal with CRISPR genome editing and can be recognized by OsTIR1 and subsequently degraded in the presence of auxin. Upon the removal of auxin, the Pol II protein would be restored. b) Western blot analyses of Pol I (RPA1), Pol II (RPB1), and Pol III (RPC1) protein levels after auxin treatment at different time points. Pol I (RPA1), Pol II (RPB1), Pol III (RPC1), Cohesin (SMC1), CTCF and TIR1(OsTIR1) protein levels were also examined here. β-actin served as a loading control. c) Lamin b1 immunofluorescence, DAPI, and GFP fluorescence signals for RNA polymerases before and after auxin treatment 6hr for Pol II, 24hr for Pol I and Pol III. Images were obtained using a 100 X objective. d) Left: Heatmap of normalized ChIP-Seq signal centered at Pol I peaks (n=3730) detected in untreated cells, and it shows marked reduction in Pol I binding in the degraded cells; middle panel for Pol II peaks (n= 40404); right panel for Pol III peaks (n= 12356). Heatmaps are ordered by descending ChIP-seq signal intensity. RNA polymerases lost chromatin binding after auxin treatment (6hr for Pol II, 24hr for Pol I and Pol III).

We next confirmed that the RPA1, RPB1, and RPC1 proteins could be depleted as an mAID-GFP fusion protein in murine embryonic stem cells. To determine the time point for Pol II loss of function, we first induced OsTIR1 expression for 24hr, added auxin, and then performed western blot analyses at different time points. Indeed, RNA polymerases could be degraded rapidly and accurately after the addition of auxin (Fig. 1b-c). RNA polymerases ChIP-Seq datasets were generated after depletion. ChIP-Seq heatmap analyses confirmed that protein depletion is efficient with the auxin-inducible degron technology (Fig. 1d). Furthermore, the global mature mRNA level measured by polyA RNA-Seq does not change dramatically after Pol II depletion for 6hr (Fig. S1b, Table S1). This rapid degradation system will allow us to specifically investigate the immediate roles of three RNA polymerases on genome structures for the first time. To determine whether the rapid depletion of RNA polymerase causes pleiotropic effects on mESCs, we compared the cellular and molecular properties of the engineered and wild-type mESCs under our experimental conditions. The cell viability, cell cycle, caspase 3/7 activities, and γH2AX level are comparable between the wild-type and the engineered cells upon treatment with doxycycline and auxin (Fig. S1c-e, S1g). These engineered cells behaved similarly to the vehicle-treated wild-type cells, indicating that the rapid depletion of RNA polymerases did not cause measurable effects on the cells at the time point that we checked and that mAID-GFP tagging did not interfere with the physiological properties of mESCs (Fig. S1c-g). A degradation time of 6hr for Pol II, 24hr for Pol I, and Pol III were chosen for the downstream Hi-C analyses because the proteins were degraded quickly. The depletion does not cause noticeable changes in the protein level of other chromatin structural regulators (i.e., SMC1 and CTCF) as well as the largest subunits of Pol I, Pol II and Pol III (Fig. 1b), although a slight destabilization of endogenously tagged proteins is observed as reported previously [40]. These results suggest that mAID-GFP fusion supports the essential functions of RNA polymerases.

### A/B Compartments and TAD structures persist after acute depletion of Pol I, Pol II, and Pol III

Previous studies on the relationships between transcription and 3D genome usually focused on Pol II [3, 42–44], here we separately investigated Pol I, Pol II, and Pol III for their roles in the 3D genome. To generate high-resolution chromatin structure data after RNA polymerases depletion, we used our recently developed BAT Hi-C (<u>B</u>ridge linker-<u>A</u>lul-<u>T</u>n5 Hi-C) [45], which could delineate chromatin conformational features such as DNA loops efficiently. Our BAT Hi-C optimized the conventional in situ Hi-C procedure by 1) fragmentation of the chromatin with the blunt four cutter restriction enzyme Alu1; 2) proximity T:A ligation with a biotinylated DNA linker; 3) isolation of chromatin to enrich chromatin-mediated interactions; and 4) construction of the library with Tn5 tagmentation (see Methods). A combination of the BAT Hi-C technique and RNA polymerase rapid degradation system will allow us to adequately investigate the relationship between specific RNA polymerases and chromatin structures.

Pol I, Pol II, and Pol III degron cells were subjected to degradation and collected for BAT Hi-C analyses. Two biological Hi-C replicates for both untreated (total reads=277 million) and Pol II degraded (total reads=300 million) cells were obtained (Table S2). The Hi-C data are reproducible (Fig. S2a) and consistent with the mESC Hi-C data quality matrix, as well as A/B compartments, published previously (Fig. S2b-c)[46]. We pooled the data and acquired a 25 kb resolution Hi-C dataset for both untreated and degraded Pol II conditions. The quality of Hi-C datasets after Pol I or Pol III degradation is similar to the Pol II degron Hi-C datasets (data not shown). Since the Hi-C detected structures are sensitive to sequencing depth, we sampled our datasets to the same sequencing depth for comparison (Table S2). Indeed, the insulation score and compartment PC1 values are comparable among the untreated Pol I, Pol II, and Pol III Hi-C datasets, and the average contact frequencies for each dataset are also similar across the genome (Fig. S2d-e).

We next sought to investigate whether perturbations occurred in A/B compartments or TADs upon the rapid depletion of RNA polymerases in mESCs. For global analyses, the A/B compartments were first delineated with Eigenvector. Then a Saddle plot displaying the compartmentalization strength indicates that the degradation of RNA polymerases does not cause apparent effects on B-B, B-A, A-B or A-A contact frequencies (Fig. 2a-c, left panel). TADs were defined with arrowhead, and we identified 1589 TADs with untreated and 1496 TADs with the Pol II degraded Hi-C dataset (Table S3). We then compared the Hi-C-detected TAD structures between untreated and degraded conditions. The TAD boundaries identified in our Hi-C dataset confirm the TAD boundaries reported previously [46]. The histogram indicates that the TAD structures are largely preserved after RNA polymerases depletion (Fig. 2a-c, middle panel). The averaged chromatin interactions around TAD structures do not reveal noticeable change (data not shown). The observed reads were resampled to equal amount divided by the expected reads (O/E) were used for calculating the averaged Hi-C signals around TAD regions. These O/E interactions would underestimate the differences but reduce the interference of experimental variations between two conditions. Consistently, the observed/expected chromatin interactions for the TAD structures do not change obviously after RNA polymerase degradation (Fig. 2a-c, right panel, Fig. S2f). For comparison, Hi-C datasets after CTCF degradation were reanalyzed with the same methods and indicated that CTCF degradation causes a significant decrease in intra-TAD interactions (Fig. 2d-e). These results indicate that the 3D genome organization persists after Pol I, Pol II, and Pol III depletion.

**Fig. 2.**
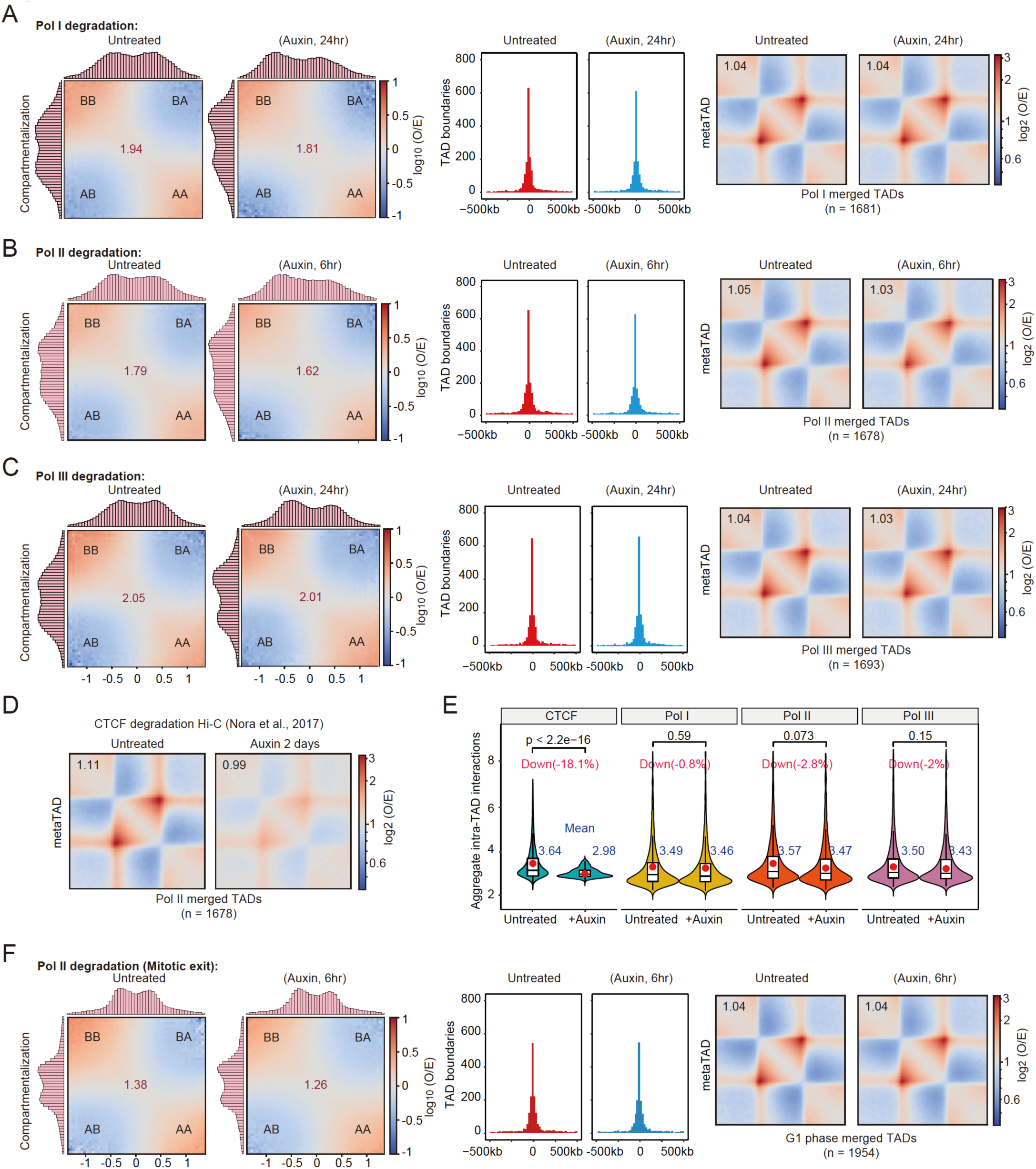
The Compartments and TADs persist after RNA polymerases depletion. a) Saddle plots representing compartmentalization strength (left), distributions of relative positions of TAD boundaries (middle), heatmap of averaged observed/expected Hi-C interactions in TAD regions (right) in untreated and Pol I degraded conditions (24hr). b) Illustrated as A, but with Pol II degradation Hi-C datasets. c) Illustrated as A, but with Pol III degradation Hi-C datasets. d) Heatmap of averaged observed/expected Hi-C interactions in TAD regions (right) in untreated and CTCF degraded conditions (2 days). e) Violin plots show quantification of the aggregated intra-TAD observed/expected contact enrichment values for various RNAP degraded conditions from BAT Hi-C data in A-C. Compared to CTCF depletion, there is no significant change in these chromatin structures after RNAP degradation for 6hr or 24hr. P values were calculated by two-sided Wilcoxon test. f) Illustrated as A, but with Pol II degradation during mitotic exit Hi-C datasets.

### Chromatin structures re-establish during mitotic exit after Pol II depletion

Our evidence suggested that RNA polymerases have little or no impact on maintenances of the 3D genome, but RNA polymerases might function during the process of 3D genome establishment. To explore this possibility, we performed Pol II degradation followed by chromatin structure analyses during mitotic exit. Previous studies have reported that TAD structures disappear in mitosis and re-establish in the early G1 phase[47]. We synchronized our cells into the M phase and simultaneously degraded Pol II and then collected both the untreated and Pol II degraded cells for Hi-C analyses during mitotic exit. The cell cycle analyses indicate that approximately 90% of cells are synchronized into M phase following previously published protocols (Fig. S2g)[48]. By comparing the cells with or without Pol II during mitotic exit, we found that degradation of Pol II during mitotic exit does not cause changes for the A/B compartments and TAD structures (Fig. 2f). These results reveal that the Pol II protein is nonessential for both the maintenance and establishment of TAD structures in mESCs.

### Pol I, Pol II and Pol III loss mildly alter local chromatin interactions

RNA polymerases are not required for global chromatin organizations; we next asked whether they play a crucial role in organizing local chromatin structures associated with active transcription. To test whether they organize local chromatin structures, we first identified Pol I, Pol II and Pol III binding hotspots or clusters with the ROSE algorithm and found mild differences in some specific regions (Fig. 3a-c). Overall, the Hi-C contact frequency decreased 2.3%, 12.6%, and 1.5% at their binding clusters, respectively for Pol I, Pol II, and Pol III (Fig. 3d), also noticed mild differences in some specific regions (Fig. 3a-c). Interestingly, Pol III binding hotspots seem to be more clustered after Pol III depletion (Fig. 3c), which is in agreement with the previously known insulator functions of tRNA elements [49–51], and is also consistent with the small but increased chromatin interactions across tRNA cluster after Pol III depletion (Fig. 3c-d). Further correlation analyses among Hi-C interaction changes and different types of functional genomic datasets were performed, and we found that Hi-C interaction changes have a better correlation with the corresponding ChIP-Seq signals (Fig. 3e). These results lead us to conclude that Pol I, Pol II, and Pol III play a modest role in structuring local chromatin interactions, and Pol II seems to contribute more. We then focused on Pol II in the following study.

**Fig. 3.**
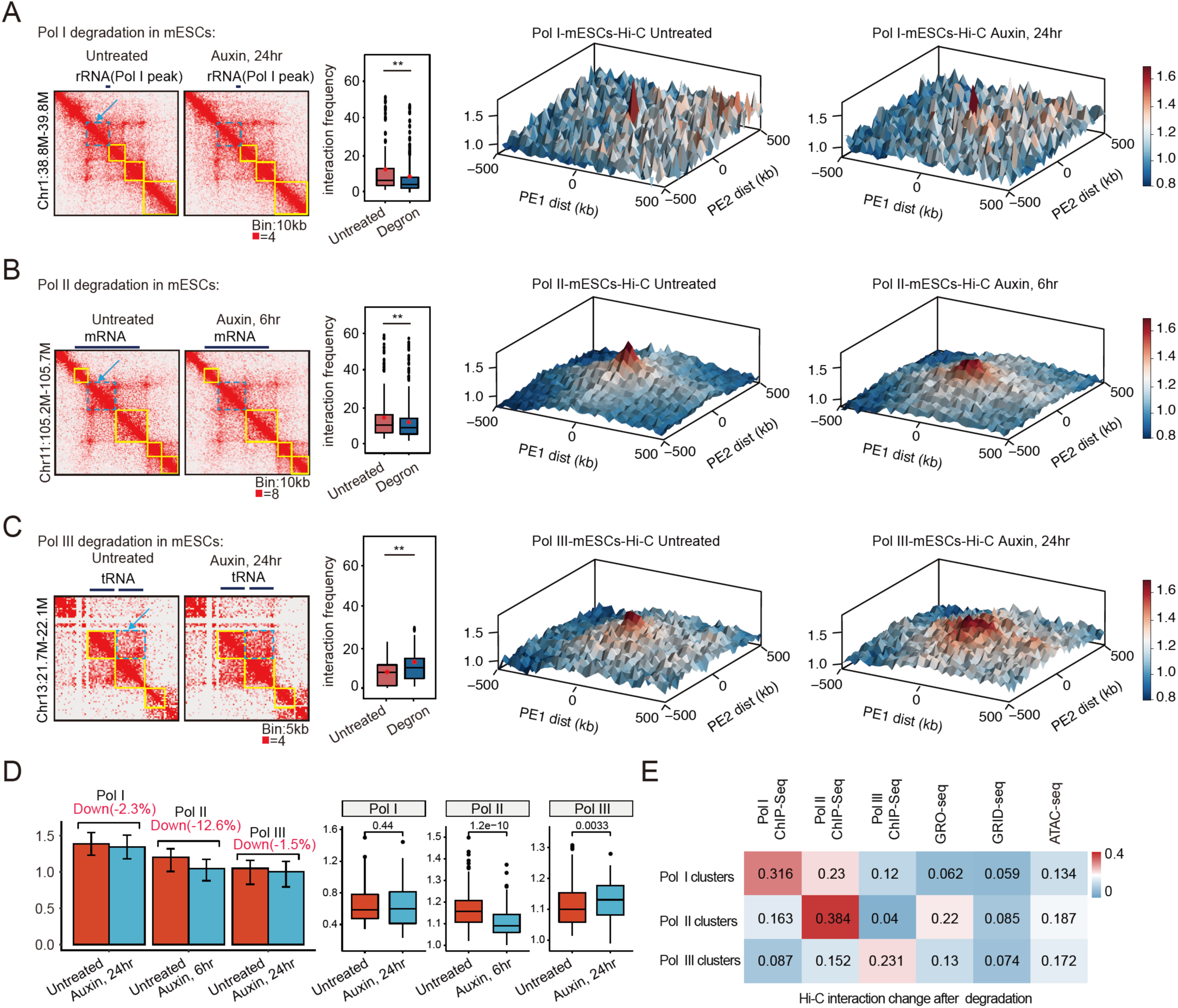
RNA polymerases depletion causes little or no changes for the local chromatin interactions. a) Hi-C contact maps for Pol I peak: 38.8-39.8 Mb region of chromosome 1. The yellow line marked regions were insulated domains, and the signals for the blue dash line marked regions were quantified and illustrated with a box plot in the left panel. Significance was determined using the Wilcoxon test (**p < 0.01). PE-SCAn analysis (10-kb resolution) depicting Hi-C contact frequency for high-density clusters of Pol I between untreated and degraded conditions were displayed in the right panel. Area shown was centered on the respective RNAP-binding sites (including 500 kb upstream and downstream). Y-axis indicates Hi-C contact frequency. b) Hi-C contact maps for mRNA (*Tlk2*) loci: 105.2-105.7 Mb region of chromosome11 at 10 kb resolution with untreated (left) and auxin-treated (right) conditions with Pol II degron cell line. The B was illustrated as A, but with Pol II degradation Hi-C datasets. c) Hi-C contact maps for tRNA cluster: 21.7-22.1 Mb region of chromosome13 at 10 kb resolution with untreated (left) and auxin-treated (right) conditions with Pol III degron cell line. The C was illustrated as A, but with Pol III degradation Hi-C datasets. d) Bar graph quantifies the reproducible highest Hi-C contacts intensity (means ± SEM) at detected Pol I, Pol II, or Pol III binding clusters (red bars) compared with the intensity of its degraded cell (blue bars) at each respective region in the left panel. Box plot displaying the change in the average contact frequency that resides within Pol I, Pol II, or Pol III binding clusters after auxin treatment in the right panel. For all box plots, center lines denote medians; box limits denote 25th–75th percentile. P values were calculated using a two-sided Wilcoxon test. e) Correlation between Hi-C contact frequency changes at Pol I clusters (top), Pol II clusters (middle), and Pol III clusters (bottom) after Pol I, Pol II, or Pol III depletion and Pol I, Pol II, and Pol III ChIP-Seq, GRO-Seq, GRID-Seq and ATAC-Seq signals in mESCs.

### Higher-resolution chromatin interaction analyses indicate that Pol II contributes no or little to organize small loop domains

The chromatin interaction analyses are highly dependent on the resolution of structures. To obtain higher-resolution information on the Pol II-dependent intra-TAD structures, we performed H3K27Ac HiChIP and Ocean-C analyses after Pol II depletion. H3K27Ac HiChIP maps chromatin loops-associated with active promoters and enhancers [52, 53] and Ocean-C, a recently developed antibody free chromosome conformation capture technique, combines FAIRE-seq with Hi-C to map hubs of open chromatin interactions [54]. Chromatin loops were identified by hichipper and HiCCUPS [55, 56] (Table S4). The histogram shows that most of the loop strength does not change much after Pol II degradation for loops identified by both algorithms with the Hi-C, HiChIP, or Ocean-C dataset (Fig. S3). Scatter plot and box plot of HiCCUPS-identified chromatin loops with all three datasets consistently show a slight decrease after Pol II depletion (Fig. 4a).

**Fig. 4.**
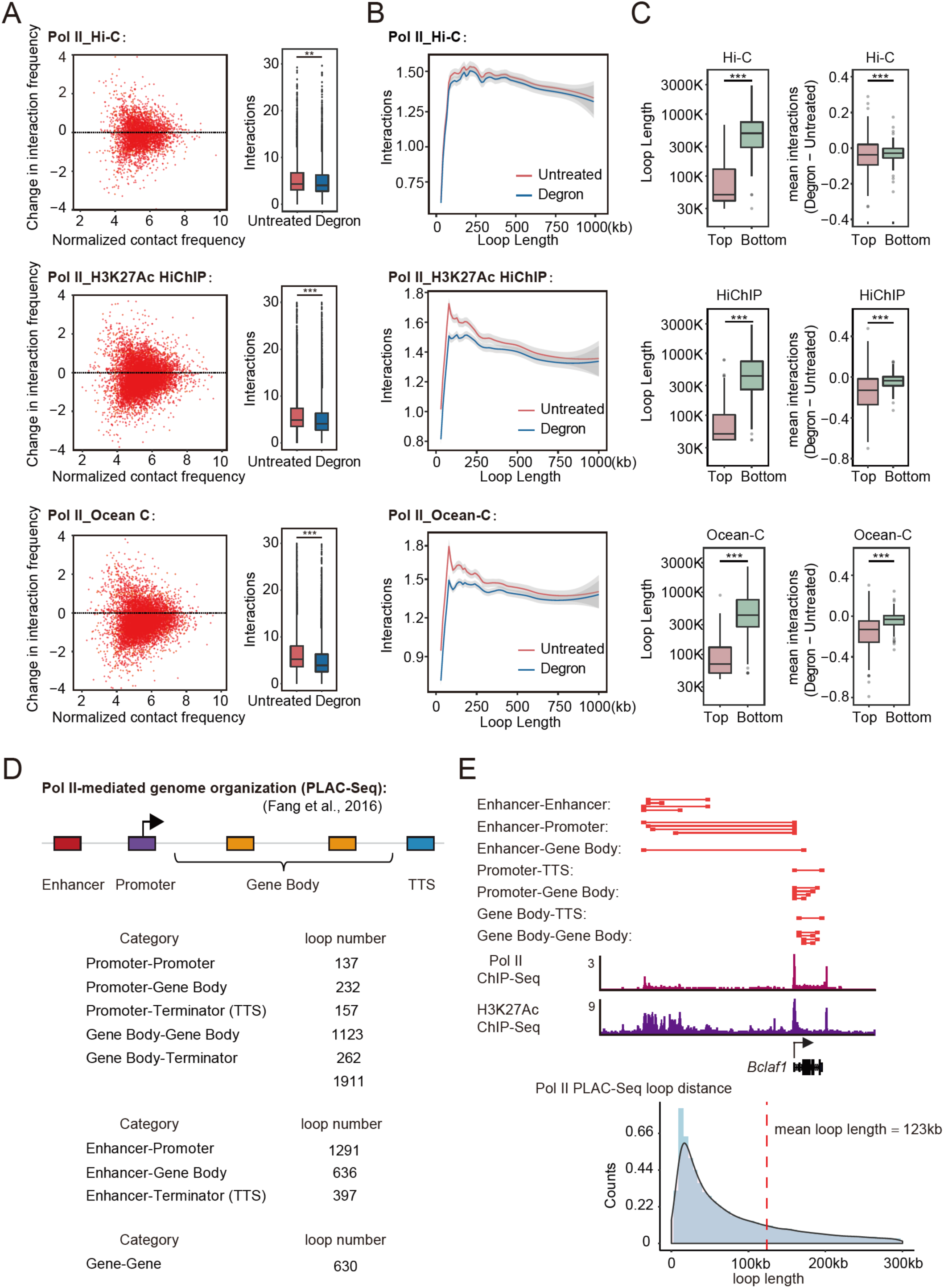
Higher resolution chromatin interaction analyses indicated that Pol II loss mildly alters actively transcribed small loop domains. a) Left: Scatter plot of the log fold change of contact frequency over normalized mean contact frequency of chromatin loops before and after Pol II degradation. Contact frequency was measured by Hi-C(top), H3K27Ac HiChIP (middle), Ocean-C (bottom). Right: Box plot of contact frequency of loops in untreated and degron cells. Significance were calculated by student t-test (***: <0.001, **: <0.01). b) Mean contact frequency ranked by the length of chromatin loops for untreated and degron conditions using the same dataset as Fig. 4A. Data are smoothed by loess regression. c) Loop length (left) and changes of mean contact frequency inside the loop domains (right) for highly transcribed (top) and lowly transcribed (bottom) chromatin loops with the same dataset as A. Significance were calculated by Kolmogorov-Smirnov test (***: <0.001). d) Global analyses for Pol II PLAC-Seq-identified chromatin loops at gene regulatory elements (enhancers, promoters, gene bodies, and terminators (TTS)). The category was listed in the left panel, and the loop counts for each category were listed in the right panel. e) The top panel showed Pol II-mediated chromatin loops (red lines). ChIP-Seq profiles for Pol II and H3K27Ac were shown at the *Bclaf1* locus (middle). A schematic of the Pol II-mediated 3D chromatin structure at the *Bclaf1* locus was shown in the right panel. The histogram of loop counts ranked by the length of loops was shown at the bottom.

We next sought to identify features of the chromatin loops that are sensitive to Pol II depletion. Previous studies reported that active transcription defined the small compartmental domains throughout Eukarya, but transcription itself could not predict 3D chromatin organization in mammals at specific loci [13]. Our analyses of chromatin structures after Pol II degradation were used to explore the existence of Pol II transcription-dependent loop domains. Loop length analyses indicate that Pol II degradation mostly affects the interaction strength for small loop domains (Fig. 4b), usually less than 250 kb, which is reminiscent of small compartmental domains in *Drosophila* [13]. Considering the current resolution of Hi-C is not sufficient to make a conclusion, we analyzed two other higher resolution datasets, HiChIP and Ocean-C, and both of them exhibited consistent trends as Hi-C did. The HiChIP and Ocean-C datasets show 14% and 15% changes more significantly than the Hi-C dataset (4%) because they preferentially capture interactions connecting open chromatin, and a similar sequencing depth would have a higher resolution than Hi-C.

If Pol II is involved in structuring small chromatin loops, we will anticipate that active and silent regions concerning Pol II transcription would behave differently after Pol II depletion. To test this idea, we ranked averaged GRO-Seq (nascent transcript) signals for loop domains and their upstream and downstream 100 kb windows in decreasing order. Indeed, the top GRO-Seq signal-associated loop domains are smaller and showed a more significant decrease in chromatin interactions than the bottom GRO-Seq signal-associated domains in the three different datasets (Fig. 4c). These results indicate that although Pol II has no functions on the global genome structures, it mildly contributes to actively transcribed small loops.

If Pol II contributes to local chromatin organization, then Proximity Ligation-Assisted ChIP-seq (PLAC-seq) analyses for Pol II should show that Pol II is preferentially associated with short-range chromatin loops [57]. We reanalyzed previously published Pol II PLAC-Seq in mESCs. The results show that Pol II-associated interactions connect promoters, gene bodies, terminators, and enhancers (Fig. 4d, Table S5). Most of these interactions are within 100kb (Fig. 4e), constrained within CTCF loop domains [58, 59]. Some Pol II-associated interactions involve chromatin clusters among genes and their potential regulatory elements, as observed in the *Bclaf1* locus (Fig. 4e). These results indicate that Pol II mediates short-range chromatin loops around gene regulatory elements and that Pol II depletion mildly affects these loops.

### Pol II bound but CTCF and Cohesin unbound loops are resistant to Pol II depletion

Our RNA polymerases degron system can degrade Pol II within 1hr (Fig. 1b), which has the unique advance to separate the impact of transcription from the presence of RNA polymerases. So, we performed BAT Hi-C analyses after Pol II depletion for 1hr and 6hr. These Hi-C datasets are highly reproducible, as we showed before (Fig. S2a, S4a). The Compartment and TAD structure analyses reproducibly show no obvious changes after Pol II depletion for 1hr and 6hr (Fig. 2b, 2f, S4b-e). Taken together, our results demonstrate that Pol II proteins are dispensable for global chromatin interactions in mESCs.

Previous transcription inhibition followed by chromatin structure analyses also exhibited no change through aggregated analyses [17, 18, 22]. These analyses did not exclude the possibility that subsets of chromatin loops might change after transcription inhibition. On the other side, most of the chromatin loops in mammals are occupied by CTCF and Cohesin, so it is better to select Pol II bound, but CTCF and Cohesin unbound loops to evaluate the contributions of Pol II in 3D genome organization. We then selected Pol II bound loops in the absence of Cohesin and found CTCF binding almost invisible at these loops (Fig. 5a). A similar classification was applied to our Pol II degron Hi-C datasets, and these loops are largely preserved after Pol II degradation for 1hr and 6hr (Fig. 5b). Then, we also identified Pol II bound but both CTCF and Cohesin unbound loops and found they also preserved after Pol II depletion (Fig. S5a). We then performed the same analyses with the previous transcription inhibition Hi-C dataset in early embryo development and activated B cell [18, 22]. The analyses consistently show that Pol II predominantly bound loops are resistant to transcription inhibition (Fig. 5b, S5a). In contrast, CTCF bound loops significantly decrease interaction strength after CTCF depletion with the same analyses pipeline (Fig. S5b).

**Fig. 5.**
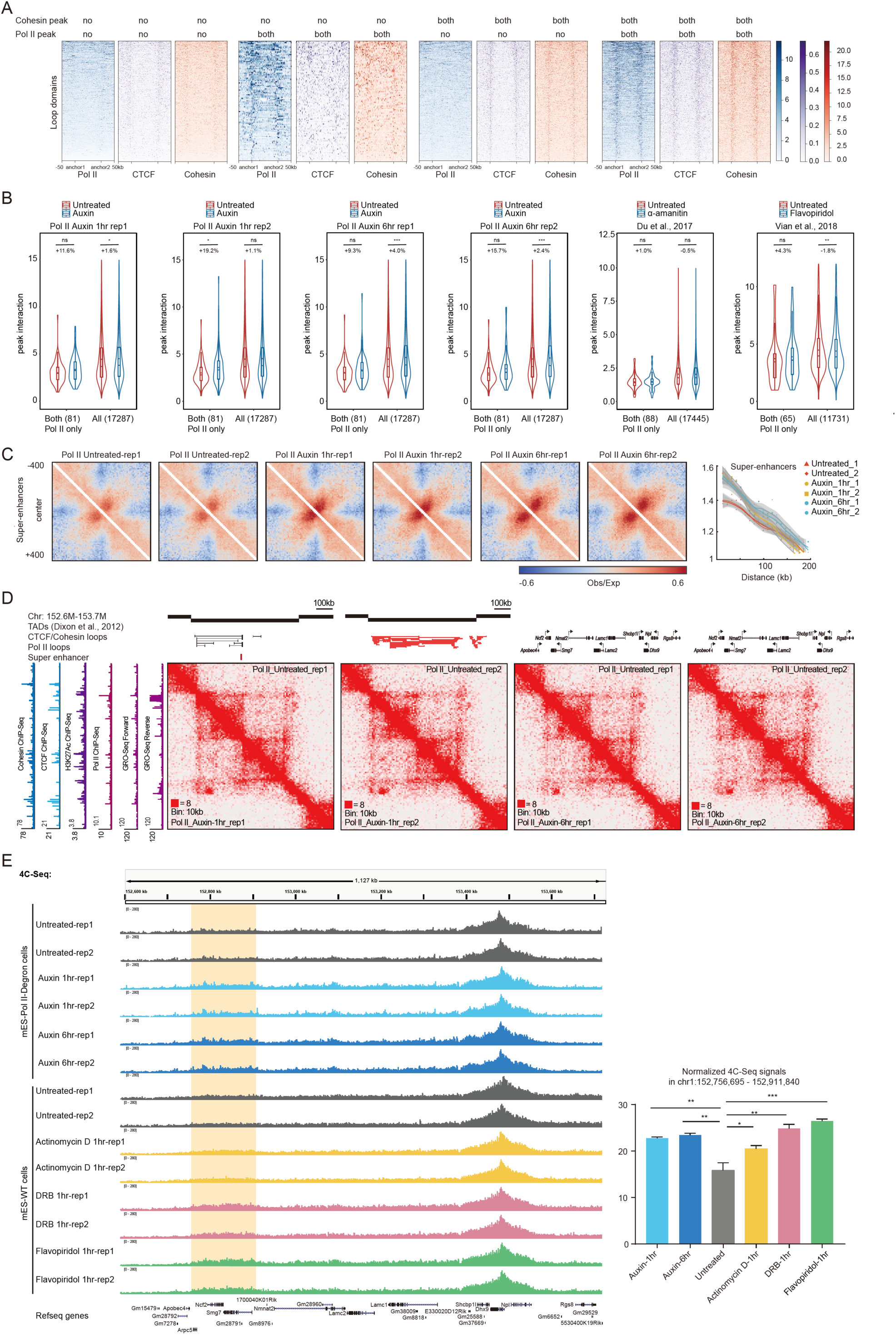
Pol II but without CTCF and Cohesin bound loops do not be affected after transcription inhibition. a) Heatmap of ChIP-Seq signal across loops classified by their overlaps of Cohesin or Pol II peaks. “Both” means both of loop anchors are overlapped with ChIP-Seq peaks, and “no” means none of the anchors overlaps with ChIP-Seq peaks. Loops are from H3K27Ac HiChIP datasets. The number of each group is: 773, 101, 2649, 787. b) Violin plots of loop peak interaction in Hi-C data of untreated and auxin(/inhibitors) conditions. Loops of Pol II bound at both anchors and Cohesin absent at both anchors are selected out and plotted with all loops. The number of loops is indicated in parentheses. The dot inside the boxplot denotes mean interaction, and the percentage of change is calculated as the difference of mean value divided by the untreated mean value. Significance were calculated by student t-test (***: <0.001, **: <0.01, *:0.05). c) Aggregate target-centered Hi-C maps reveal that Pol II degradation slightly increased of Super-enhancer (SE) stripes formation. Pile-up maps were plotted by the 10kb resolution contact matrix and normalized to distance, with a positive signal in red and negative in blue. In the right panel, signal decaying curve of the SE stripes. The x-axis shows the enrichment of SE stripes, and the y-axis shows the distance to the target center up to 200 kb. d) Hi-C contact maps for *Dhx9*: 152.6-153.7 Mb region of chromosome 1 at 10 kb resolution with untreated and auxin-treated Pol II degron cell line. The Cohesin, CTCF, H3K27Ac ChIP-Seq signals, and GRO-Seq signals were displayed on the left. The black bars indicate as TAD structures. The black lines indicate as CTCF/Cohesin loops, and the red lines indicate as Pol II-associated loops. e) The 4C analyses at the *Dhx9* locus after Pol II degradation and transcription inhibitions were shown. Transcription was inhibited with flavopiridol, actinomycin D, or DRB for 1hr. The orange background regions indicated the 4C enriched region, and its quantitative analysis was shown at the right. The untreated signals indicate the average signals of untreated conditions for mES-WT and mES-Pol II degron cells.

Super-enhancers loci are usually associated with extremely high levels of transcription [60, 61]. Therefore, we examined these regions for the changes in chromatin interactions. The target-centered maps indicate that the super-enhancer regions mildly increase chromatin interactions after Pol II degradation at both 1hr and 6hr (Fig. 5c). Specifically, the interactions around the key pluripotency gene *Esrrb* and housekeeping gene *Dhx9* adjacent to super-enhancer-associated loop domains are also increased (Fig. 5d, S5c). Moreover, this observation could be further independently validated by 4C-Seq analyses after Pol II depletion (Fig. 5e). We then inhibited transcription with actinomycin D, DRB, or flavopiridol and observed a consistent increase trend in the interaction frequency (Fig. 5e). Recent live-cell imaging analyses indicate that Pol II transcription restrains the dynamics of chromatin [62]. This explains our observation that Pol II degradation causes a slight increase in chromatin interactions of super-enhancer regions. These results further suggest that Pol II is not directly involved in chromatin organization but restrain the dynamics of chromatin, underlying a mechanism for the increase of chromatin interactions after Pol II depletion.

### Immediate depletion of Pol II does not perturb Cohesin chromatin binding, but more prolonged depletion does

If Pol II indeed is nonessential in 3D genome organization, how to explain the previous finding that transcription-mediated Cohesin chromatin binding and transcription elongation-mediated chromatin structures? We speculated that Pol II might indirectly regulate Cohesin. Further taking advantage of the Pol II degron system, we can degrade Pol II at different time points, especially at 1hr time point (Fig. 1b). The single locus (Fig. 6a) and meta-gene analyses (Fig. 6b-c) for ATAC-Seq signals and Cohesin (Smc1) ChIP-Seq show that these signals do not change after Pol II degron for 1hr but decrease for 6hr. These results implicate that Pol II degradation may indirectly change the 3D chromatin interaction. Inhibiting transcription for more extended time points, many potential secondary effects may emerge (decrease of short half-life transcripts, noncoding RNAs, chromatin accessibility and stress of transcription inhibition, etc.). Our data here underpin a model that transcription inhibition does not cause immediate changes of chromatin interactions while indirectly decreases chromatin interactions at Pol II clusters potentially through decreasing chromatin accessibility and Cohesin binding on chromatin.

**Fig. 6.**
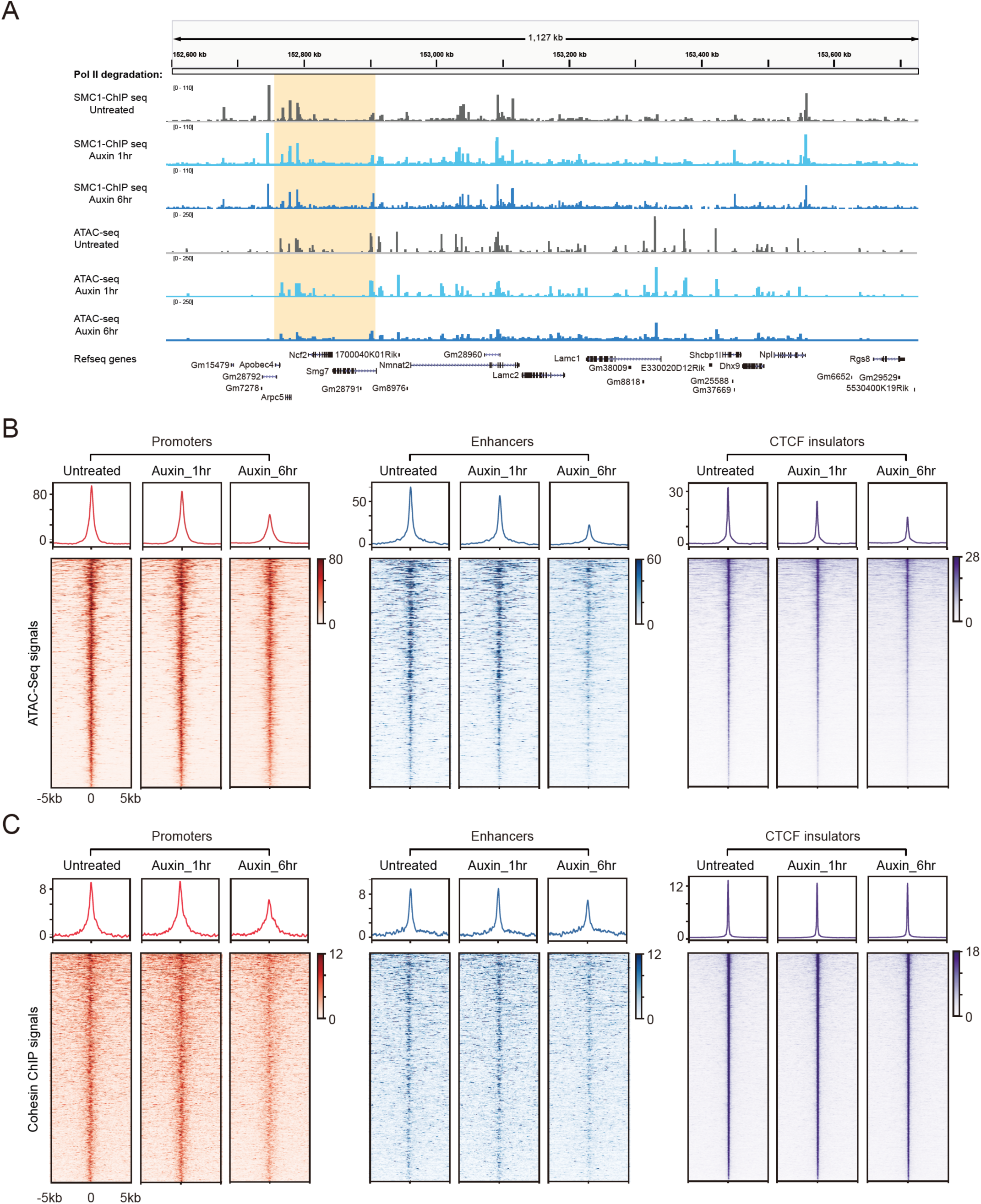
Pol II depletion for a longer time significantly affects chromatin accessibility and Cohesin occupancy. a) ChIP-Seq profiles for Cohesin (SMC1) and ATAC-seq signals at the *Dhx9* locus before and after Pol II degradation for 1hr and 6hr were shown. The orange background regions correspond to the 4C enriched region in Fig 5e. b) Heatmap illustrating ATAC-seq signals centered at all promoters, enhancers, and CTCF-binding insulators (± 5kb) before (untreated) and after auxin treatment (1hr or 6hr). Those regions were ranked by their level of chromatin accessibility (from high to low) for each catalog in mESCs. c) Same analysis as in Figure 6b but focused on Cohesin ChIP-seq binding.

## Discussion

The relationship between transcription and the 3D genome organization is one of the most fundamental questions in modern molecular biology. Here we provided evidence: 1) specific transcription inhibition by degradation of Pol I, Pol II and Pol III result in little or no changes for global 3D chromatin structures as assayed by Hi-C (Fig. 2a-c, 2f); 2) select high abundance Pol I, Pol II and Pol III binding sites-associated chromatin interactions and found they still persist after degradation (Fig. 3a-c); 3) generate higher resolution chromatin interaction maps and reveal that transcription inhibition mildly alters small loop domains (Fig. 4b-c); 4) separate the impact of transcription from the presence of RNA polymerase by the degradation of Pol II within 1hr and show that Pol II protein is dispensable for global chromatin organization (Fig. S4b-d); 5) identify Pol II dominant loops in the absence of CTCF and Cohesin and found they are mostly resistant to transcription inhibition (Fig. 5b, S5a). This evidence collectively demonstrates RNA polymerases play a marginal role in the 3D genome organization. Besides, time-course experiments show that more prolonged depletion causes a decrease of Cohesin binding, but not for the short time treatment (Fig. 6b-c), suggesting that RNA polymerases can regulate the 3D chromatin landscape indirectly. RNA polymerases are core enzymes for transcription that are constitutively expressed in all cell types, so 3D chromatin organization independent of transcription is likely to be general in mammalian cells.

Mounting evidence has shown a strong correlation between transcription and 3D genome structures [12, 63, 64], while their relationship is still controversial before our study. Transcription inhibition experiments suggest the persistence of chromatin structures after transcription inhibition [17, 18], but they either performed only aggregated analyses or based on a single resolution chromatin interaction map. So, they cannot exclude the possibility that transcription might play a predominant role in organizing specific types of chromatin structures or with higher resolution interaction analyses. None of the previous studies have thoroughly investigated the transcription roles in 3D chromatin interaction, as we did in this study. Our conclusion here, transcription inhibition, causes no or little effects on 3D chromatin structures, is consistent with many previous studies and is essential for the field. On the other side, many studies have shown that transcription inhibition affects 3D genome structures and Cohesin (Busslinger et al., 2017; Heinz et al., 2018), which seems to conflict with our conclusion. Our time-course experiments clearly show that more prolonged depletion of Pol II decreases chromatin accessibility and Cohesin binding, but not at the shorter time point of Pol II degradation, suggests that transcription can regulate 3D genome structures indirectly. Our study perfectly explains the previously confusing results: transcription inhibition usually does not affect 3D genome structures, but the high dosages, long time treatment, or more sensitive biological systems may cause chromatin structure change via indirectly affecting Cohesin.

There are many genome architectural proteins (such as CTCF, YY1, Znf143, RNA polymerases, RNA binding proteins and mediators, etc.) [24, 25, 65–71], and noncoding RNAs have been implicated in organizing 3D genome structures, so the 3D genome may likely organize via a combinatory of many factors. In this study, we show that the Pol I, Pol II, and Pol III binding hotspots, their chromatin interactions, and Pol II only bound loops are resistant to transcription inhibition. These results argue against this combinatory model for protein factors, but we cannot exclude a possibility that nuclear noncoding RNAs may organize 3D genome structures, which can also be independent of transcription activities.

We show that transcription may indirectly affect 3D genome structures via Cohesin chromatin binding, and there are many other possibilities. For example, a previous imaging study showed that Pol II transcription constrains the dynamics of chromatin in the nucleus. After transcription inhibition, the highly transcribed regions move fasters in the nucleus, interact more with some regions, and less with the other regions. Since transcription is vital for all genes, it is also likely that transcription affects short half-life chromatin structural proteins.

Previous studies showed that structural factors such as CTCF and Cohesin-mediated chromatin interactions create frameworks for loop domains and constrain high-frequency chromatin interactions within them [2, 3, 72, 73]. CTCF and Cohesin depletion destroy most chromatin structures in mammalian cells, while it has little effects on transcription [24, 26]. Here we depleted the core subunits of Pol I, Pol II, and Pol III in mESCs and found little changes for chromatin structures. This data suggests that transcription and 3D chromatin structures are largely uncoupled in the nucleus of mESCs. The previous signaling induced and heat shock followed chromatin structure analyses suggested a pre-existing model for the 3D genome [74–76]. Our evidence also supports the pre-existing model, for which the mechanisms could be connected to genome sequences or epigenetic modifications.

## Conclusions

Our study provides the first comprehensive analyses of the roles of Pol I, Pol II, and Pol III proteins in 3D chromatin organization in mESCs. We demonstrate that RNA polymerases play a minor role in organizing the 3D genome, and we propose that transcription does not regulate the 3D genome directly but is able to regulate the 3D genome indirectly. This explains the confusing effects on the 3D genome after transcription inhibition because it is difficult to separate the direct or indirect effects of transcription inhibition. Our study also implicates that transcription and 3D chromatin organizations are largely uncoupled in the nucleus. Since transcription is nonessential for the 3D genome, further studies of genetics, epigenetics features of 3D genome, and noncoding RNAs may reveal new insights for the 3D genome organization.

## Materials and methods

Details see supplementary information file.

## Additional files

**Fig. S1.**
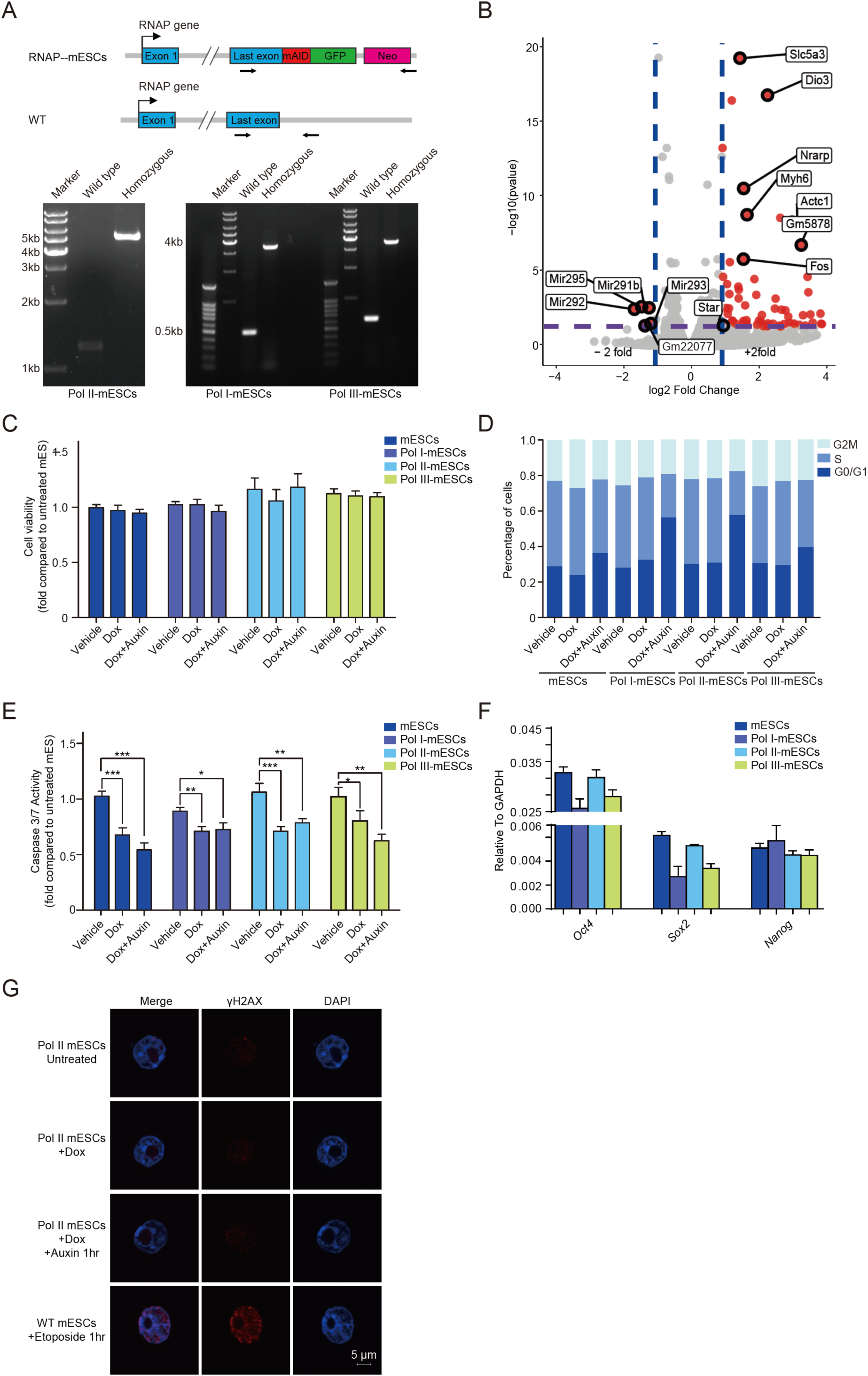
Characterization of the RNAP degron cell lines and quality control for Hi-C. **a.** Design of CRISPR knock-in for RNAP-mAID-GFP fusions in mESCs. Primer for RNAP-mAID-GFP genotyping indicated as arrowhead (top). Genotyping for Pol I-mAID-GFP, Pol II-mAID-GFP, Pol III-mAID-GFP mESCs were shown in the bottom panel. **b.** Volcano plot displaying poly-A RNA-Seq signals for untreated and Pol II depleted (auxin treated for 6hr) mESCs. **c-e.** Cell viability (c), cell cycle (d), and caspase 3/7 (e) analyses for wild type, Pol I, Pol II, Pol III mAID-GFP fusion mESCs under the treatment of vehicle doxycycline, doxycycline and auxin. **a. f.** RT-qPCR analyses for *Oct4*, *Sox2*, and *Nanog* with wildtype, Pol I (RPA1), Pol II (RPB1), and Pol III (RPC1) mAID-GFP fusion mESCs. **b. g.** Immunofluorescent staining for γH2AX (red; left) and DAPI (blue; right) signals after 1hr individual drug treatment were shown. Images were obtained using a 100 X oil objective.

**Fig. S2.**
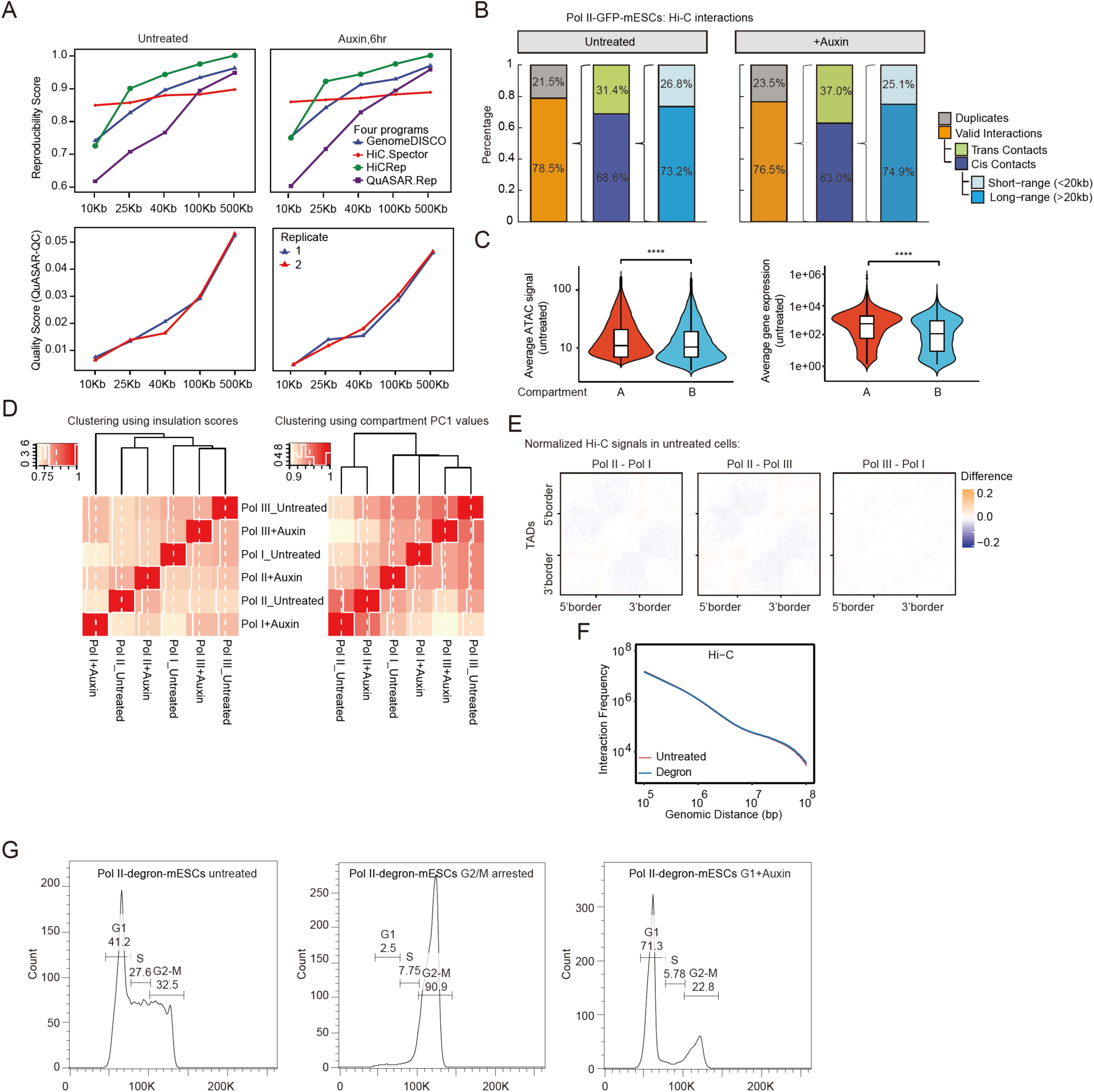
Quality controls for Pol I, Pol II, and Pol III degradation Hi-C datasets. **a.** Reproducibility analysis of BAT Hi-C replicates. Reproducibility scores were calculated by four independent algorithms, including GenomeDISCO, QuASAR, HiC.Spector and HiC-Rep. The charts plot the reproducibility score (y-axis) at different resolutions (x-axis). All methods reported at least 60% of the reproducibility rate from 10kb to 500kb resolutions. The Quality Score was measured by QuASAR. **b.** The alignment statistics of BAT Hi-C assay for the untreated (total reads=277 million) and Pol II degraded (total reads=300 million) samples. Typically, the BAT Hi-C library contains over 75 % of unique pairs with an optimal amount of PCR cycles. About 70 % of valid pairs are cis contacts that consist of 26.8 % short-range interactions (<20 kb) and 73.2 % long-range interactions (>20 kb), while 30% of pairs are inter-chromosomal contacts. **c.** Compartment A identified in this study had higher ATAC-Seq signals (left panel) and gene expression (right panel) compared with compartment B, indicating that our compartment identification works. Significance was determined using the Wilcoxon test (****p < 0.0001). **d.** Left panel: Hierarchical clustering of the compartment A/B scores (PC1 values) between samples. Right panel: Hierarchical clustering of the insulation scores between samples. **e.** Average observed/expected Hi-C interactions within TADs of Pol II-mAID mESCs untreated subtracted Pol I-mAID-mESCs (left) or Pol III-mAID-mESCs (middle) untreated ones. The averaged observed/expected Hi-C interactions within TADs of Pol III-mESCs untreated subtracted Pol I-mESCs untreated signals were shown in the right panel. **f.** Averaged Hi-C interaction frequency distribution along genomic distance in untreated and degron conditions. **g.** Cell cycle analyses for the mitotic exit experiment with Pol II-degron cells.

**Fig. S3.**
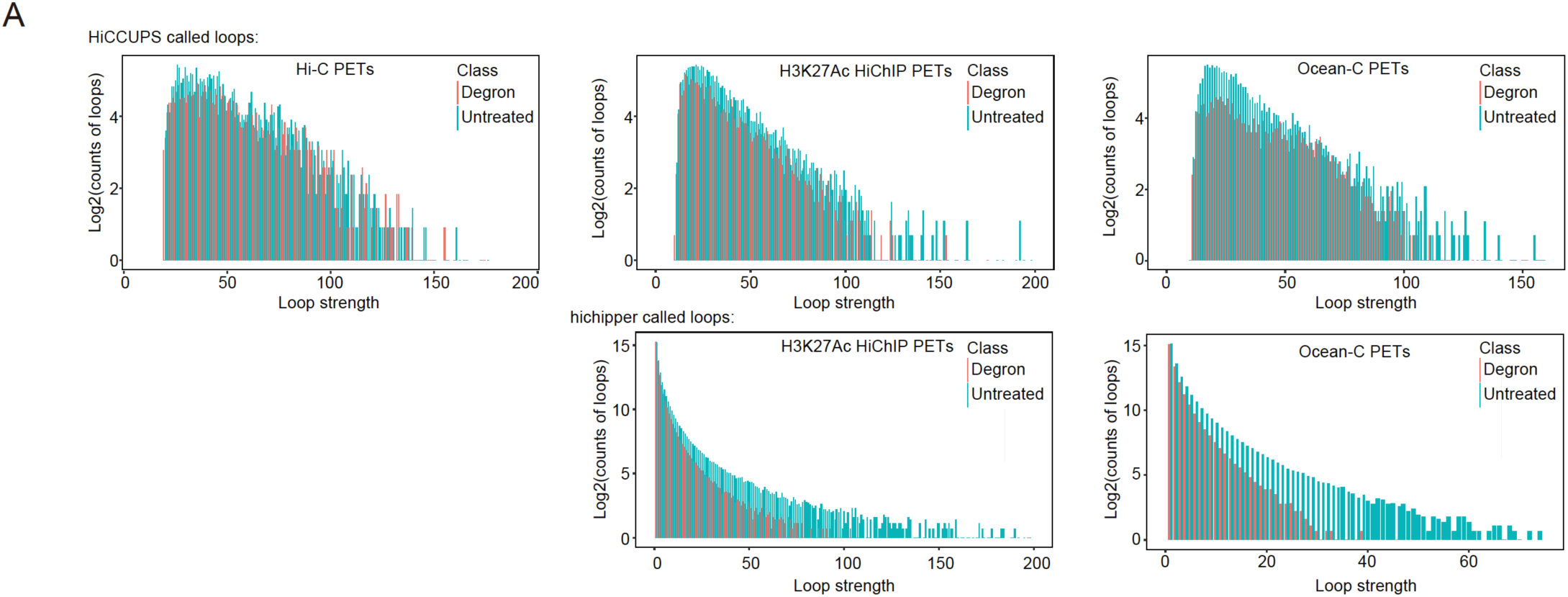
Loop strength analyses of higher resolution chromatin interactions after Pol II depletion. **a.** Loop strength distribution analyses of Hi-C (left), H3K27Ac HiChIP (middle) and Ocean-C (right). The x-axis denotes the loop strength, and the y-axis denotes the log counts of loops with different strengths. Top: loops were called by HiCCUPS; bottom: loops were called by hichipper. Red bars indicate degron conditions, and blue bars indicated untreated conditions.

**Fig. S4.**
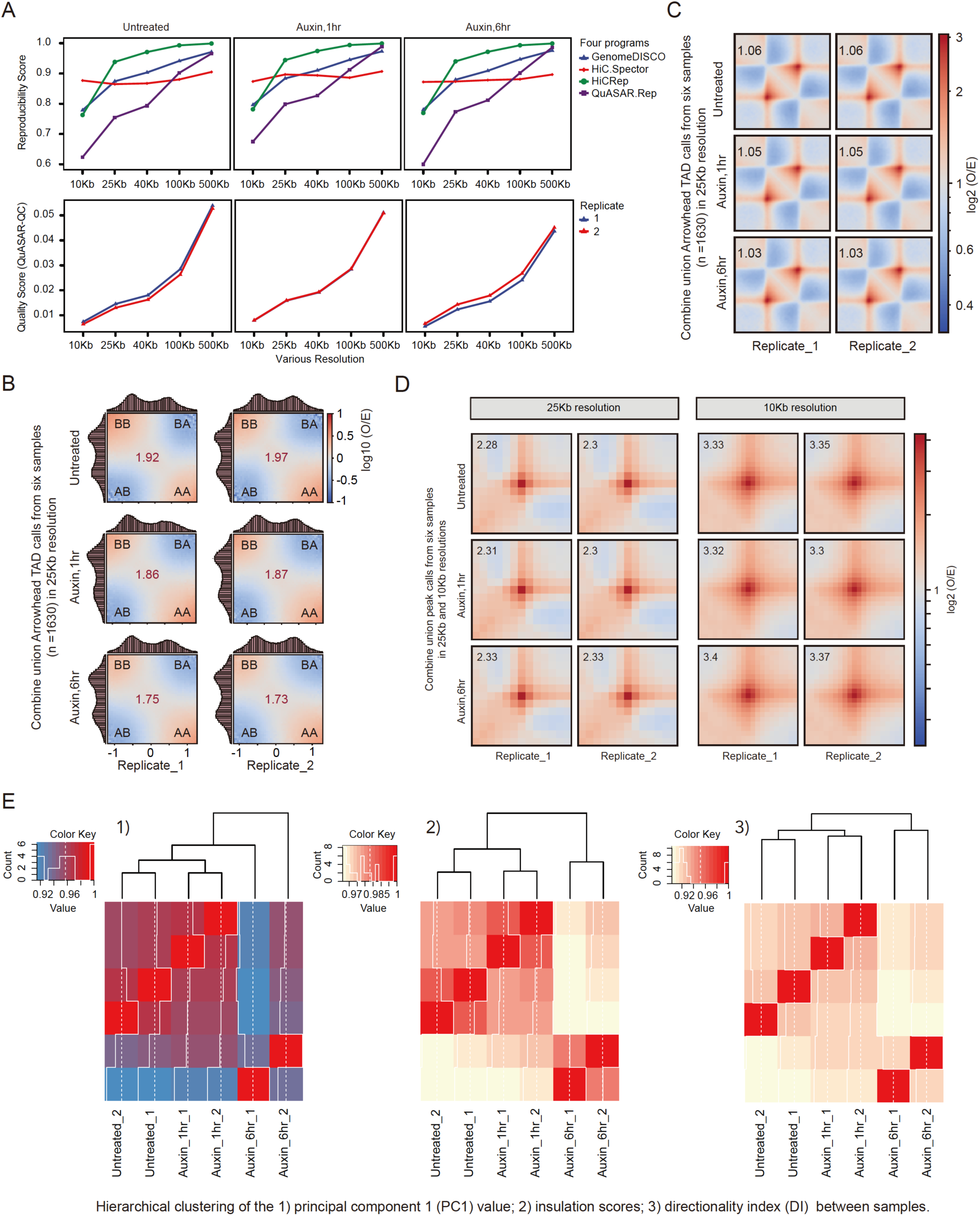
Quality controls for Pol II degradation time course Hi-C datasets. **a.** Same analysis as in Figure S2A, but with the time course of Pol II degron Hi-C datasets. **b.** Saddle plots representing compartmentalization strength with the time course of Pol II degron Hi-C datasets. **c.** Heatmap of averaged observed/expected Hi-C interactions with the time course of Pol II degron Hi-C datasets. **d.** Aggregated peak analysis (APA) of the union loop calls on merged and two biological replicates from untreated and Pol II degraded cells. Bin size, 10kb and 25kb. Numbers in the top left corner of each heatmap indicate average loop strength: log2(obs/exp). Colour is shown in log-scale and shows enrichment of interactions. **e.** Hierarchical clustering of the 1) compartment A/B scores (PC1 values), 2) insulation scores, 3) directionality index (DI) between samples.

**Fig. S5.**
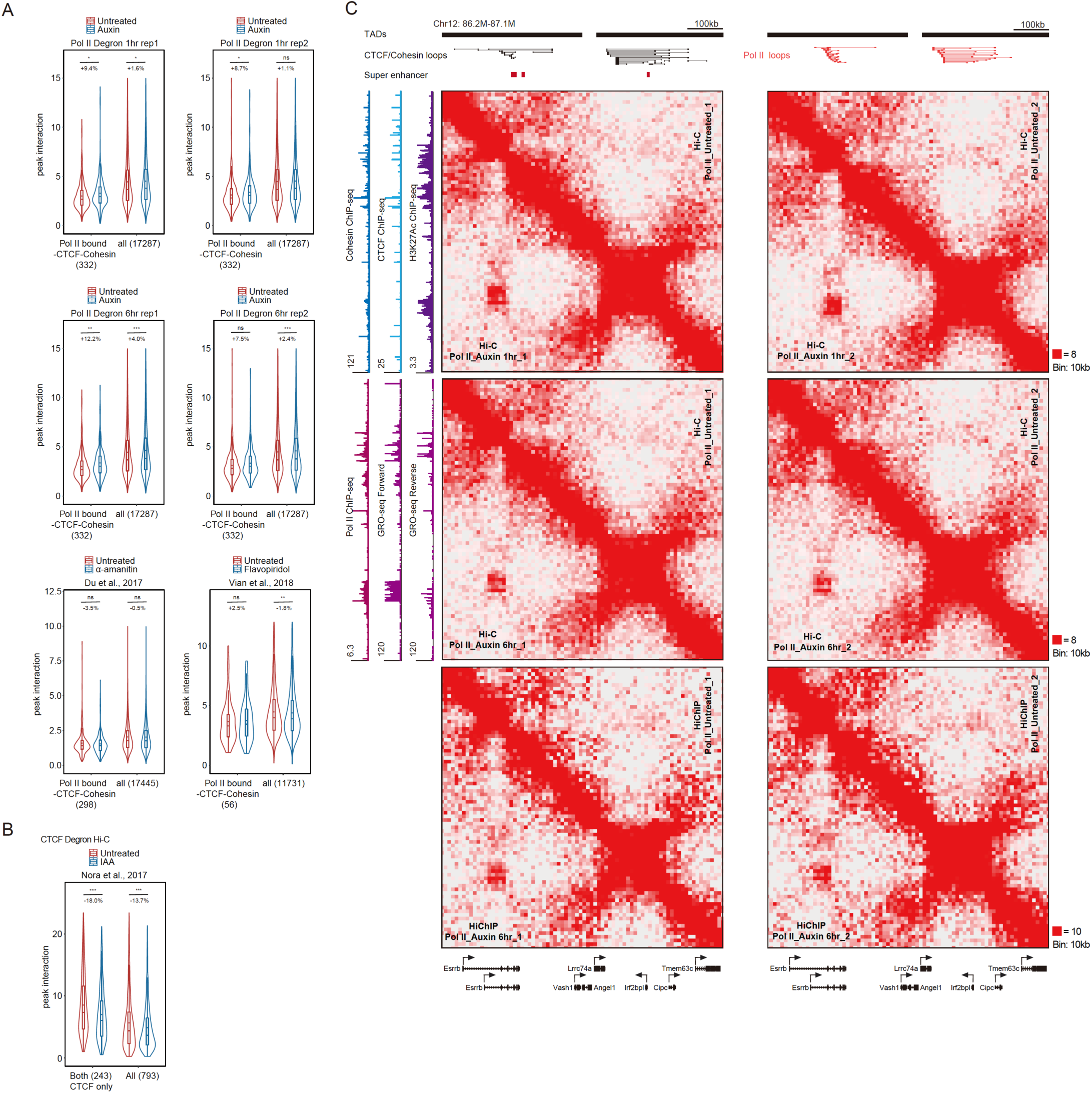
Examples of super-enhancer regions mildly gain chromatin interactions across loop domains after Pol II depletion. **a.** Violin plots of loop peak interaction in Hi-C data of untreated and auxin(/inhibitors) conditions. Loops of Pol II bound at either anchor but both CTCF and Cohesin unbound loops are selected out and plotted with all loops. The number of loops is indicated in parentheses. The dot inside the boxplot denotes mean interaction, and the percentage of change is calculated as the difference of mean value divided by the untreated mean value. Significance were calculated by student t-test (***: <0.001, **: <0.01, *:0.05). **b.** Violin plots of loop peak interaction in CTCF Hi-C data of untreated and degron conditions. Loops of CTCF bound at both anchors are selected out and plotted with all loops. b is illustrated similar to a. **c.** Hi-C (upper) and H3K27Ac HiChIP (Bottom) contact maps for *Esrrb*: 86.2-87.1Mb region of chromosome 2 at 10kb resolution with untreated and auxin-treated Pol II degron cell lines. The Cohesin, CTCF, H3K27Ac ChIP-Seq signals, and GRO-Seq signals were displayed on the left. The black bars indicate as TAD structures. The black lines indicate CTCF/Cohesin loops, and the red lines indicate as Pol II-associated loops.

**Additional file 6: Table S1.** RNA-seq RPKM values in Pol II Untreated and Degron cells. **Table S2.** Summary statistics of the Hi-C, HiChIP, and Ocean-C data. **Table S3.** Hi-C identified TADs (contact domains). **Table S4.** HiCCUPS identified loop domains. **Table S5.** Pol II PLAC-seq high confidence interactions identified using the Origami pipeline. **Table S6.** RNAP ChIP-Seq Peaks. **Table S7.** PCR primer sequences used in this study. **Table S8.** List of datasets used in this study.

## Abbreviations

TADs: Topologically associating dominas
ChIP: chromatin immunoprecipitation
CTCF: CCCTC-binding factor
DAPI: 4’,6-diamidino-2-phenylindole
DRB: 5,6-Dichloro-1-beta-Ribo-furanosyl Benzimidazole
GRO-Seq: Global nuclear run-on sequencing
Ocean-C: open chromatin enrichment and network Hi-C
GRID-Seq: global RNA interactions with DNA by deep sequencing
PLAC-Seq: proximity ligation assisted ChIP-seq
RNAP: RNA polymerase

## Authors’ contributions

X.J. conceived and supervised the project. Y.P.J. generated degron mESCs, performed all of sequencing and qPCR experiments with the initial help from R.Z., D.W.J., Y.Q.F.. H.N.Z. performed 4C-Seq and γH2AX staining experiments and analyses with help from Y.P.J. and B.Y.L.. Y.J.L. performed the characterization analysis for degron cells, Y.H. performed IF analyses for localization. J.H., K.H.L. and B.Y.L. performed all the bioinformatics. All authors contributed to data analysis and data interpretation. X.J. wrote the manuscript with input from Y.P.J., J.H., K.H.L., B.Y.L., H.N.Z. and help from other authors.

## Acknowledgements

We thank Drs. Xiang-dong Fu, Richard Young, and the members of the Ji laboratory for their discussions. We thank Lumeng Jia for advice on Ocean-C experiments, Dr. Jiazhi Hu, Jianhang Yin for advice on 4C-seq experiment, Dr. Wei Xie, Du zhenghai for transcription inhibition Hi-C datasets, Dr. Chunyang Wang and Wenhong Jiang for initial data analysis. We thank the National Center for Protein Sciences at Peking University for assistance with imaging and flow cytometry. This work was supported by funds from the Ministry of Science and Technology of China and the National Natural Science Foundation of China (Grants 2017YFA0506600 and 31871309), the Young Thousand Talents Program of China, the grants from the Peking-Tsinghua Center for Life Sciences, and the Key Laboratory of Cell Proliferation and Differentiation of the Ministry of Education at Peking University School of Life Sciences to X. Ji. J. H. was supported by a grant from the China Postdoctoral Science Foundation.

